# Three-dimensional imaging reveals distinct muscle-vessel-nerve relationships during dorsal and ventral muscle splitting in the chicken forelimb

**DOI:** 10.64898/2026.09.16.749688

**Authors:** Yuta Takase, Sayaka Tojima

**Affiliations:** Bioorganic Research Institute, Suntory Foundation for Life Sciences, Kyoto, Japan; Applied Biology, Kyoto Institute of Technology, Kyoto, Japan

**Keywords:** chicken embryo, forelimb development, muscle splitting, vasculature, peripheral nerves, tissue clearing, three-dimensional imaging

## Abstract

During vertebrate limb development, muscles, blood vessels, and nerves undergo coordinated morphogenesis. Limb muscles form through progressive splitting of the dorsal and ventral muscle masses, but the three-dimensional relationships between muscle splitting and surrounding tissues remain incompletely understood. Although blood vessels have been implicated in muscle splitting, whether this association is shared among different muscle masses is unclear. Here, we established a three-dimensional imaging approach for chicken embryonic forelimbs by combining fluorescent vascular labeling, whole-mount immunostaining, tissue clearing, and confocal microscopy. We compared three hydrophilic clearing methods, Sca*l*eS, CUBIC, and SeeDB2, for compatibility with fluorescent ink, LCA-lectin, and PKH26 vascular labeling. SeeDB2 preserved all three vascular signals. We then developed a modified SeeDB2 protocol incorporating blood decolorization and used it to examine muscle, vessel, and nerve relationships from E5.0 to E6.0. In the ventral muscle mass, multiple vascular extensions were observed around the splitting region at E5.5. In contrast, no prominent vascular extension was detected around the splitting region of the dorsal muscle mass despite ongoing splitting at the same stage. Nerve projections also showed distinct spatial patterns between the dorsal and ventral muscle masses. These findings indicate that vascular extension is not a universal feature of limb muscle splitting and suggest that individual splitting events occur within distinct local tissue environments. Our three-dimensional approach provides new morphological insight into the coordinated development of muscle, vasculature, and peripheral nerves in the embryonic limb.

## INTRODUCTION

During vertebrate limb development, multiple tissues, including the skeleton, muscles, tendons, blood vessels, and nerves, undergo morphogenesis in parallel, and their proper spatial organization and integration are essential for the formation of a functional musculoskeletal system. Interactions among developing muscles, tendons, and skeletal elements play important roles in this process, making the limb a widely used model for studying the coordinated morphogenesis of multiple tissues (Huang, 2017). Blood vessels and peripheral nerves also develop in close association within the embryonic limb, and a high degree of three-dimensional spatial correspondence between the two has been reported in the avian forelimb (Bates et al., 2002). Muscle and nerve development likewise proceed concurrently; in the chicken embryonic forelimb, nerve fibers begin to invade the limb bud around E5, while dorsal and ventral muscle masses form and subsequently subdivide into individual muscles (Luxey et al., 2020). Thus, although muscles, blood vessels, and nerves develop within the same spatiotemporal environment, their local three-dimensional relationships during limb muscle morphogenesis remain incompletely understood.

Limb muscles originate from muscle progenitor cells derived from the dermomyotome of the somites. These cells migrate into the limb bud, where they form dorsal and ventral muscle masses that are subsequently subdivided in a stepwise manner to generate individual muscles (Scaal & Marcelle, 2018; Smith-Paredes et al., 2022). This latter process, referred to as muscle splitting, is a key morphogenetic event that contributes to establishing the final shape and spatial arrangement of individual muscles. Recent three-dimensional comparative analyses across multiple amniote species have shown that early muscle mass cleavage/splitting patterns are highly conserved among lineages, highlighting the developmental significance of splitting patterns for understanding muscle homologies (Smith-Paredes et al., 2022). Migration of muscle progenitor cells into the limb is regulated by several signaling pathways, including HGF/c-MET, SDF1/CXCR4, and ephrin/EPH signaling, and local signals from the limb mesenchyme also contribute to muscle patterning (Scaal & Marcelle, 2018). However, how established muscle masses are spatially subdivided into individual muscles within their local tissue environment remains incompletely understood.

Muscle splitting has been proposed to be influenced, at least in part, by the developing vasculature. In the chicken embryonic forelimb, endothelial cells have been reported to be positioned across future cleavage sites between E5 and E5.5, before clear separation of the ventral muscle mass occurs (Tozer et al., 2007). Moreover, experimentally increasing ectopic vascularization suppresses muscle formation, whereas local inhibition of vascular development results in muscle fusion, supporting a functional role for blood vessels in muscle splitting (Tozer et al., 2007). However, it remains unclear whether this relationship between vascular patterning and muscle splitting is shared among different muscle masses or splitting events. Peripheral nerves also project into the muscle masses during this developmental period, and close spatial relationships between nerves and early muscle splitting have been suggested in other amniotes (Smith-Paredes et al., 2022). Nevertheless, three-dimensional analyses directly comparing the spatial relationships of splitting sites with both blood vessels and nerves in the chicken embryonic forelimb remain limited.

To address this question, it is important to examine developing limb muscles, blood vessels, and nerves while preserving their complex three-dimensional organization. Traditionally, the spatial relationships between blood vessels and muscles in the chicken embryonic forelimb have been analyzed mainly using histological sections. However, the developing vascular network undergoes dynamic remodeling through branching and regression, while nerves and blood vessels follow complex three-dimensional trajectories, limiting comprehensive assessment of inter-tissue relationships from two-dimensional sections alone (Bates et al., 2002). More recently, combinations of whole-mount immunostaining, tissue clearing, and light-sheet fluorescence microscopy have enabled three-dimensional analyses of muscle and nerve architecture in the chicken embryonic limb, as well as muscle splitting patterns across amniotes (Luxey et al., 2020; Smith-Paredes et al., 2022). Tissue-clearing methods, however, differ substantially in principle, including hydrophobic, hydrophilic, and hydrogel-based approaches, and vary not only in clearing efficiency but also in their ability to preserve fluorescent proteins and chemical fluorescent tracers (Tian et al., 2021; Ueda et al., 2020). Therefore, when blood vessels are visualized by introducing fluorescent labeling reagents into the vasculature, clearing conditions must be selected to preserve the labeling signal.

To address these issues, we first compared tissue-clearing methods for their compatibility with several fluorescent vascular labels introduced into the chicken embryonic vasculature and identified conditions that preserved the vascular labeling signals. We then developed a modified SeeDB2 protocol incorporating blood decolorization to improve tissue transparency and combined this method with fluorescent vascular labeling and whole-mount immunostaining to analyze muscle splitting in E5-E6 chicken embryonic forelimbs. We found that vascular extensions were present around the splitting region of the ventral muscle mass, whereas no comparable vascular extensions were observed in the dorsal muscle mass despite ongoing splitting. Nerves also displayed distinct spatial patterns in the ventral and dorsal muscle masses. These findings indicate that vascular extension is not a universal feature of muscle splitting and suggest that distinct local tissue environments may contribute to individual splitting events in different regions of the developing limb.

## MATERIALS AND METHODS

### Chicken embryos

Fertilized chicken eggs were purchased commercially from Yamagishi Poultry Farms (Wakayama, Japan). Embryos were staged according to the Hamburger and Hamilton (HH) staging system (Hamburger & Hamilton, 1951) or by embryonic day (E). All procedures involving chicken embryos, including vascular labeling and fixation, were performed at Kyoto Institute of Technology and were approved by the Animal Experimentation Committee of Kyoto Institute of Technology (approval no. 100237) in accordance with institutional guidelines.

### Visualization of blood vessels

*In vivo* visualization of blood vessels was performed by intravascular infusion of highlighter ink (PILOT Spotliter; Takase et al., 2013), Lens culinaris agglutinin lectin (LCA-lectin; Jilani et al., 2003), or PKH26 Cell Linker (Sigma-Aldrich, MINI26). For highlighter ink infusion, 0.5-1 μl of diluted highlighter ink (PILOT Spotliter; 1:5 in phosphate-buffered saline (PBS)) was injected into chicken embryos through the heart or yolk sac vasculature using a micropipette prepared from a glass capillary (Narishige, GD-1) with a vertical micropipette puller (Narishige, PC-10). For lectin infusion, 0.5-1 μl of 0.5 mg/ml Fluorescein-labeled LCA-lectin (Vector Laboratories, FL-1041-5) was injected in the same manner. For PKH26 infusion, 0.5-1 μl of PKH26 solution (1:2 dilution in Diluent C) was injected through the heart or the right dorsal aorta. After injection, the embryos were incubated at 38.5°C for an additional 5 min before harvesting. Fluorescence images were acquired using a Leica MZ10 F stereomicroscope equipped with a DS-Ri1 camera (Nikon) or an FV3000 confocal microscope (Evident).

### Sample preparation and tissue clearing

Chicken embryos were fixed overnight in 4% paraformaldehyde (PFA; Nacalai Tesque) in PBS at 4°C and then washed three times in PBS for 5 min each. Tissue clearing was subsequently performed using three different methods: Sca*l*eS, CUBIC, and SeeDB2. Each method was performed basically according to the original reports (Hama et al., 2015; Ke & Imai, 2018; Susaki et al., 2015). For the Sca*l*eS method, samples were sequentially immersed in Sca*l*eS0 [20% (w/v) sorbitol (Nacalai Tesque), 5% (w/v) glycerol (Nacalai Tesque), 1 mM methyl-β-cyclodextrin (Nacalai Tesque), 1 mM γ-cyclodextrin (Nacalai Tesque), 1% (w/v) N-acetyl-L-hydroxyproline (Nacalai Tesque), and 3% (v/v) DMSO (Nacalai Tesque) in PBS], Sca/eS1 [20% (w/v) sorbitol, 10% (w/v) glycerol, 4 M urea (Nacalai Tesque), and 0.2% (w/v) Triton X-100 (Nacalai Tesque) in dH2O], Sca*l*eS2 [27% (w/v) sorbitol, 2.7 M urea, 0.1% (w/v) Triton X-100, and 8.3% (v/v) DMSO in dH2O], and Sca*l*eS3 [36.4% (w/v) sorbitol, 2.7 M urea, and 9.1% (v/v) DMSO in dH2O] at 4°C. The samples were then washed in PBS at 4°C for 1 h and finally incubated in Sca*l*eS4 [40% (w/v) sorbitol, 10% (w/v) glycerol, 4 M urea, 0.2% (w/v) Triton X-100, and 20% (v/v) DMSO in dH2O] at room temperature. For the CUBIC method, samples were incubated in CUBIC-1A solution [10% (w/v) Triton X-100, 5% (w/v) N,N,N,N’-tetrakis(2-hydroxyethyl)ethylenediamine (2-HP ethylenediamine) (Tokyo Chemical Industry), 10% (w/v) urea, and 25 mM NaCl (Nacalai Tesque) in dH2O] at 37°C for 2-3 h to decolorize the blood. After decolorization, the samples were washed in PBS for 1 h and then cleared in CUBIC-2 solution [25% (w/v) urea, 50% (w/v) sucrose (Nacalai Tesque), and 10% (w/v) triethanolamine (Tokyo Chemical Industry) in dH2O] at room temperature. For the SeeDB2 method, samples were first immersed in 2% (w/v) saponin (Nacalai Tesque) in PBS and then sequentially transferred to 33% (v/v) Omnipaque 350 (GE HealthCare)/2% (w/v) saponin, 67% (v/v) Omnipaque 350/2% (w/v) saponin, and finally 100% Omnipaque 350 at room temperature. For the modified SeeDB2 method, blood decolorization was performed by incubation in saponin/CUBIC-1A solution [2% (w/v) saponin, 5% (w/v) 2-HP ethylenediamine, 10% (w/v) urea, and 25 mM NaCl in dH2O] at 37°C for 2 h before replacement with Omnipaque 350.

### Immunohistochemistry

PFA-fixed chicken embryonic forelimbs were subjected to blood decolorization using the modified SeeDB2 method described above. The samples were then washed three times in 0.2% Triton X-100 in PBS (PBST) for 30 min each. After blocking with 1% bovine serum albumin (BSA; Sigma-Aldrich) in PBST for 1 h, the samples were incubated overnight at 4°C with the following primary antibodies: eFluor 660-conjugated anti-beta 3 tubulin (TUBB3) mouse monoclonal antibody (1:250; Invitrogen, 50-4510-80) and anti-myosin 4 mouse monoclonal antibody (1:250; Invitrogen, 14-6503-82), which had been labeled with HiLyte Fluor 555 using an Ab-10 Rapid HiLyte Fluor 555 Labeling Kit (Dojindo, LK35). The samples were then washed three times in PBST containing Hoechst 33342 (1:1000; Nacalai Tesque, 19172-51) for 1 h each. After washing, the samples were sequentially immersed in Omnipaque 350 according to the SeeDB2 protocol.

### Image acquisition and analysis

Fluorescence images of chicken forelimbs were acquired using FV3000 or FV4000 confocal microscopes (Evident), each equipped with a 10× objective lens. Samples were placed in a glass-bottom dish (IWAKI) filled with Omnipaque 350 and imaged under the objective lens. The image size for each optical section was set to 512 × 512 pixels with a pixel size of 1.23 μm × 1.23 μm and a z-step of 10 μm. Hoechst 33342, fluorescent yellow ink, HiLyte Fluor 555, and eFluor 660 were excited at 405, 488, 561, and 640 nm, respectively. Image processing and three-dimensional reconstruction were performed using ImageJ software (National Institutes of Health).

## RESULTS

### SeeDB2 preserves signals from multiple fluorescent vascular labels

Prior to three-dimensional visualization of muscles, blood vessels, and nerves in chicken embryonic forelimbs, we evaluated tissue-clearing methods suitable for combination with fluorescent vascular labeling (Fig. 1). Blood vessels in E2.0 (HH14) chicken embryos were labeled using three fluorescent vascular labels—fluorescent ink, LCA-lectin, and PKH26—and the fluorescence signals were compared before and after clearing with Sca*l*eS, CUBIC, or SeeDB2 (Fig. 1A). All three clearing methods increased tissue transparency compared with uncleared embryos (Fig. 1B). However, preservation of the vascular labeling signals differed among the clearing methods.

**Figure 1.**
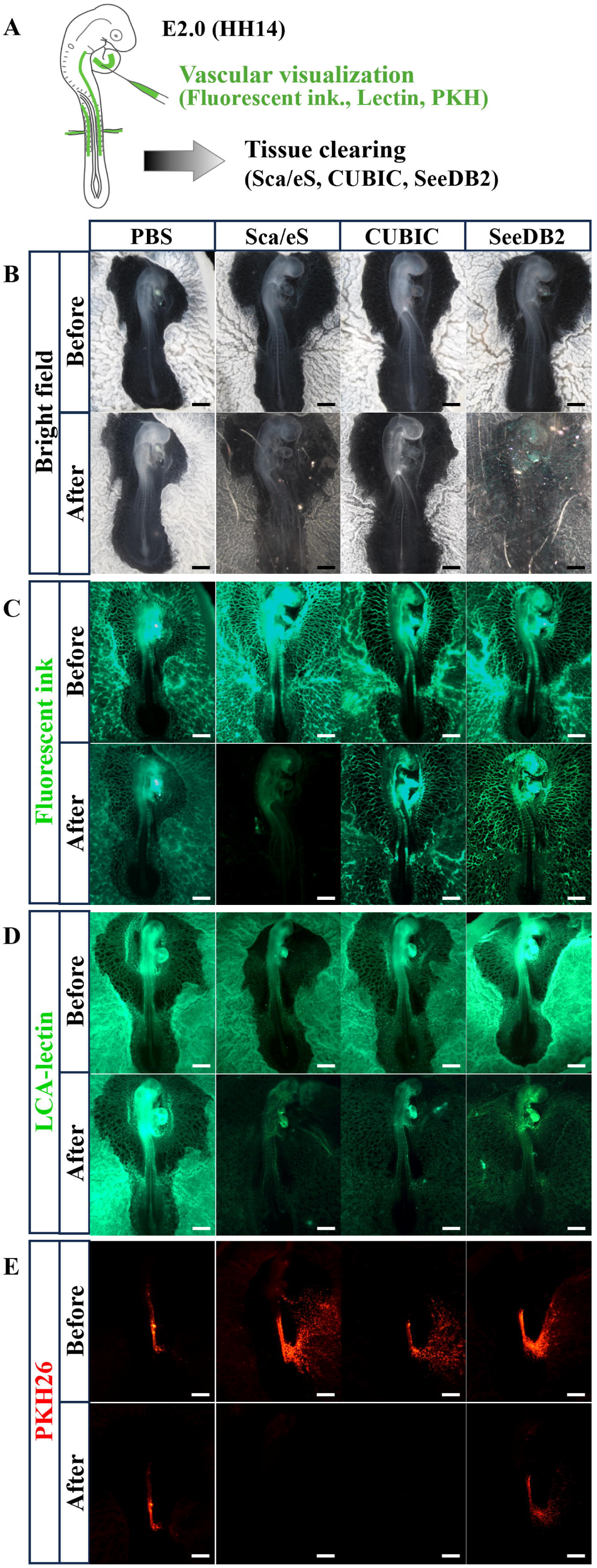
Evaluation of tissue-clearing methods compatible with fluorescent vascular labeling. (A) Schematic overview of the experimental procedure. Three fluorescent vascular labels—fluorescent ink, LCA-lectin, and PKH26—were introduced into the vasculature of E2 chicken embryos. Fluorescent ink and LCA-lectin were injected into the heart, whereas PKH26 was injected through the heart or right dorsal aorta. Embryos were then fixed with 4% paraformaldehyde (PFA) and subjected to one of three tissue-clearing methods: Sca*l*eS, CUBIC, or SeeDB2. Fluorescence signals were compared before and after clearing. (B) Bright field images before and after tissue clearing. All three clearing methods increased tissue transparency. (C) Fluorescent ink signals before and after tissue clearing. Signal attenuation was observed after Sca*l*eS treatment. (D) LCA-lectin signals before and after tissue clearing. No marked signal attenuation was observed with any of the three clearing methods. (E) PKH26 signals before and after tissue clearing. Signal attenuation was observed after Sca*l*eS and CUBIC treatment. Scale bars, 1 mm.

For vascular labeling with fluorescent ink, the signal was retained after CUBIC and SeeDB2 treatment, whereas a marked reduction in fluorescence was observed after Sca*l*eS treatment (Fig. 1C). For LCA-lectin labeling, no marked reduction in fluorescence was observed after any of the three clearing methods (Fig. 1D). In contrast, for vascular labeling with the fluorescent lipid tracer PKH26, the signal was retained after SeeDB2 treatment but was markedly reduced after Sca*l*eS and CUBIC treatment (Fig. 1E). These results indicated that SeeDB2 was compatible with all three fluorescent vascular labeling approaches examined in this study. We therefore selected SeeDB2 as the tissue-clearing method for subsequent visualization of blood vessels in chicken embryos.

### Modified SeeDB2 incorporating blood decolorization improves tissue transparency while preserving PKH26 signals

Based on the results shown in Fig. 1, we next applied SeeDB2 clearing to E6 (HH28) chicken embryonic forelimbs. After SeeDB2 treatment, residual blood remained dark within the forelimb tissue and limited tissue transparency (Fig. 2A). In contrast, when SeeDB2 clearing was performed after CUBIC treatment, which includes a blood decolorization step (Tainaka et al., 2014), residual blood was reduced and tissue transparency was improved compared with SeeDB2 treatment alone (Fig. 2B). However, as shown in Fig. 1, CUBIC treatment markedly reduced PKH26 vascular labeling signals. We therefore modified the SeeDB2 protocol to achieve blood decolorization while preserving PKH26 fluorescence.

**Figure 2.**
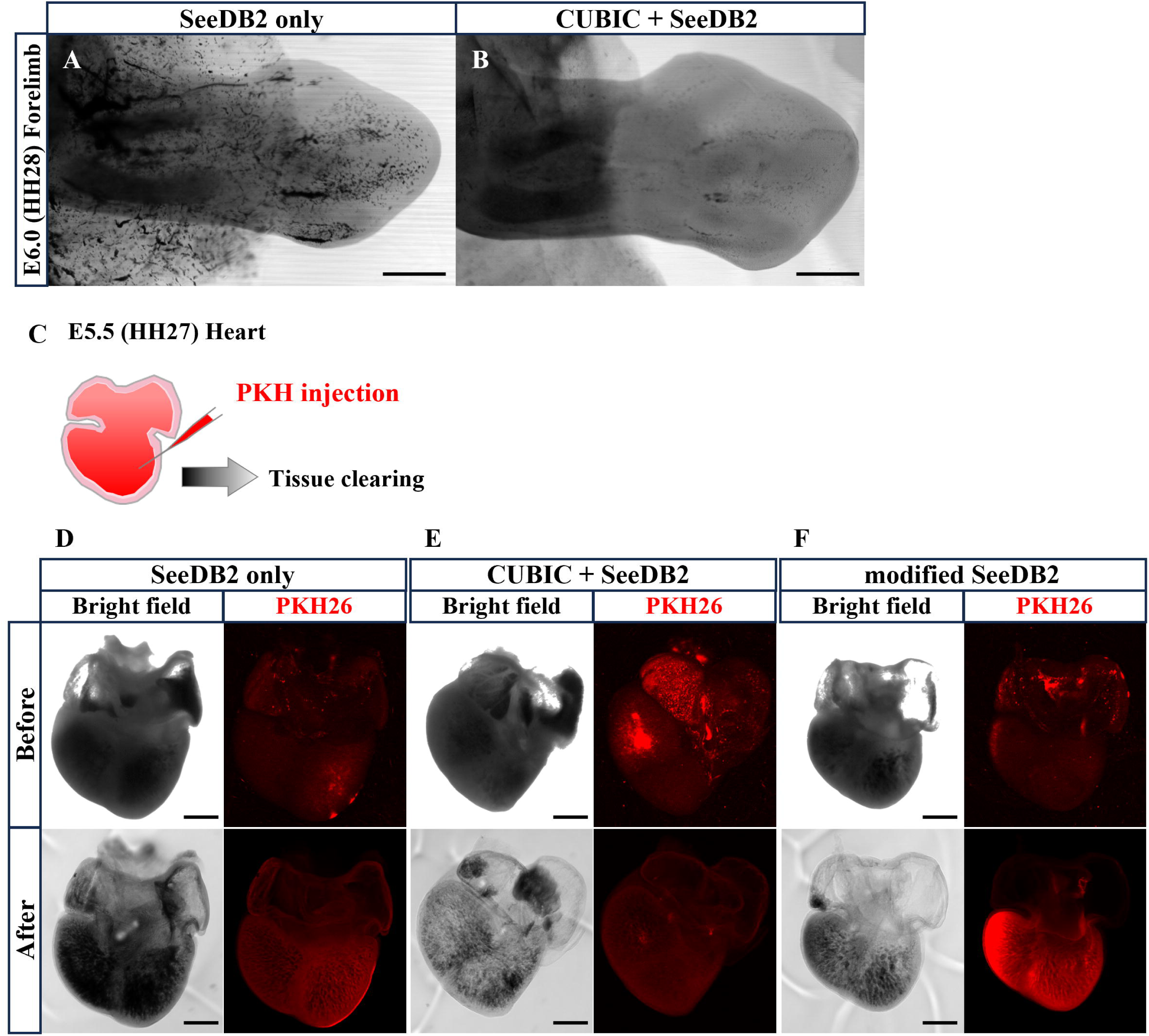
Improved tissue transparency using a modified SeeDB2 protocol incorporating blood decolorization. (A) Bright field image of an E6 chicken forelimb after SeeDB2 treatment alone. Residual blood remained within the tissue and limited tissue transparency. (B) Bright field image of an E6 chicken forelimb after CUBIC treatment followed by SeeDB2 clearing. Blood decolorization reduced residual blood and improved tissue transparency compared with SeeDB2 treatment alone. (C) Schematic overview of the experiment. PKH26 was injected into the heart of E5.5 (HH27) chicken embryos, followed by tissue clearing. (D) Bright field images and PKH26 signals before and after SeeDB2 treatment alone. PKH26 signals were preserved after clearing, whereas residual blood remained in the tissue. (E) Bright field images and PKH26 signals before and after CUBIC treatment followed by SeeDB2 clearing. Blood decolorization markedly improved tissue transparency, whereas PKH26 signals were attenuated. (F) Bright field images and PKH26 signals before and after modified SeeDB2 treatment incorporating blood decolorization with saponin. Tissue transparency was improved while PKH26 signals were preserved, allowing clearer visualization of the vasculature after clearing. Scale bars, 500 μm.

Because CUBIC-1A contains 10% Triton X-100, we hypothesized that this detergent treatment might contribute to the reduction in PKH26 fluorescence. We therefore compared three conditions using E5.5 (HH27) chicken embryos in which PKH26 had been injected into the heart: (1) SeeDB2 treatment alone, (2) SeeDB2 treatment following blood decolorization with CUBIC, and (3) SeeDB2 treatment following blood decolorization using a modified CUBIC-1A solution in which 10% Triton X-100 was replaced with 2% saponin (Fig. 2C). With SeeDB2 treatment alone, PKH26 fluorescence was preserved after clearing, but residual blood limited tissue transparency and hindered clear visualization of the heart (Fig. 2D). When CUBIC treatment was followed by SeeDB2 clearing, blood decolorization markedly improved tissue transparency, whereas PKH26 fluorescence was substantially reduced (Fig. 2E). In contrast, blood decolorization using the saponin-based treatment followed by SeeDB2 clearing improved tissue transparency while preserving PKH26 fluorescence, resulting in clearer visualization of the heart after clearing (Fig. 2F). Based on these results, we used this modified SeeDB2 protocol for subsequent three-dimensional visualization of muscles, blood vessels, and nerves in chicken embryonic forelimbs and for analysis of their spatial relationships during muscle splitting.

### Vascular extension is associated with the splitting region of the ventral muscle mass

We first focused on the ventral muscle mass of the chicken embryonic forelimb, in which vascular extension associated with muscle splitting has previously been reported (Tozer et al., 2007), and analyzed the spatial relationships among muscles, blood vessels, and nerves from E5.0 (HH26) to E6.0 (HH28) (Fig. 3A–D). Before splitting, at E5.0 (HH26), the ventral muscle mass appeared as a single, thin muscle mass (Fig. 3E’, F’). At this stage, no prominent vascular extension into the muscle mass was observed (Fig. 3E, E’), whereas some nerve projections into the muscle mass were already evident (Fig. 3F, F’). At E5.5 (HH27), when splitting was in progress, the ventral muscle mass still appeared as a continuous mass, but a putative splitting region was apparent near its center (white arrowheads in Fig. 3G’, H’). Multiple vascular extensions were observed around this region (Fig. 3G, G’). In contrast, no prominent nerve projection was observed around the splitting region; instead, nerves projected toward regions that would later form the ventro-anterior and ventro-posterior muscle masses (Fig. 3H, H’). After splitting, at E6.0 (HH28), the ventral muscle mass had separated into ventro-anterior and ventro-posterior muscle masses (Fig. 3I’, J’). No blood vessels were observed between the two muscle masses, and no prominent vascular extension into either muscle mass was evident (Fig. 3I, I’). In contrast, nerve projections were observed in both the ventro-anterior and ventro-posterior muscle masses (Fig. 3J, J’).

**Figure 3.**
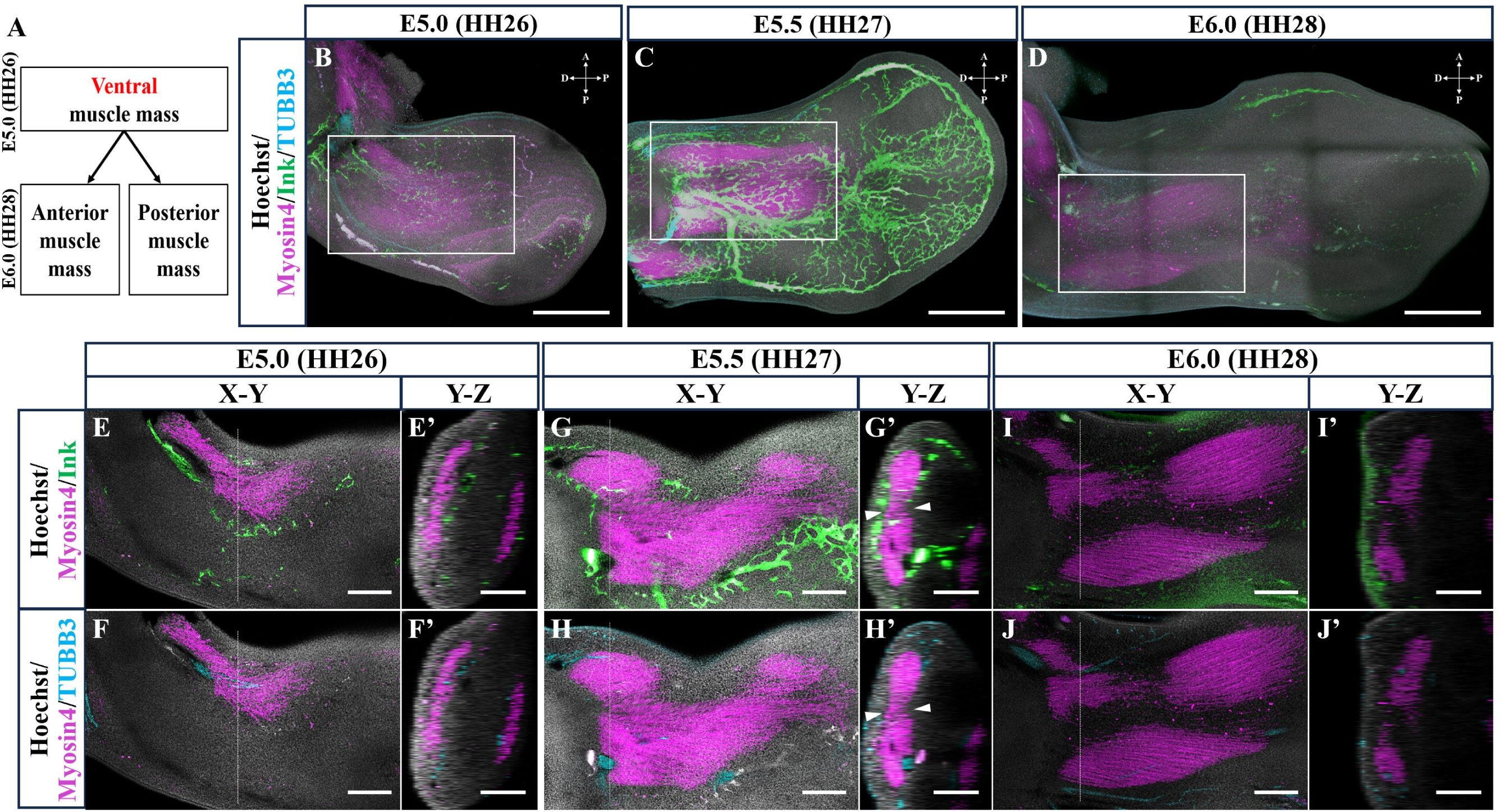
Spatial relationships among muscle, blood vessels, and nerves during splitting of the ventral muscle mass. (A) Schematic illustration of ventral muscle mass splitting from E5.0 (HH26) to E6.0 (HH28). (B–D) Muscles (Myosin4), blood vessels (Fluorescent ink; Ink), and nerves (TUBB3) in chicken embryonic forelimbs before splitting (E5.0 [HH26]; B), during splitting (E5.5 [HH27]; C), and after splitting (E6.0 [HH28]; D). White boxes indicate regions shown at higher magnification in E–J’. (E–J’) Higher-magnification X–Y images and corresponding Y–Z views at the positions indicated by dashed lines. At E5.0, no prominent vascular extension into the ventral muscle mass was observed (E, E’), whereas nerve projections were already present (F, F’). At E5.5, multiple vascular extensions were observed around the putative splitting region (white arrowheads; G, G’), while no prominent nerve projection was detected around this region; nerves instead projected toward regions that would later form the ventro-anterior and ventro-posterior muscle masses (H, H’). At E6.0, no blood vessels were observed between the separated ventro-anterior and ventro-posterior muscle masses, and no prominent vascular extension into either mass was evident (I, I’), whereas nerve projections were present in both muscle masses (J, J’). Nuclei, muscles, blood vessels, and nerves were visualized with Hoechst, anti-Myosin4, fluorescent ink, and anti-TUBB3, respectively. Scale bars: 500 μm in B–D; 200 μm in E–J and E’–J’.

Together, these observations indicate that blood vessels and nerves show distinct spatial patterns during splitting of the ventral muscle mass, with vascular extension becoming prominent around the splitting region at E5.5.

### Dorsal muscle splitting occurs without prominent vascular extension around the splitting region

We next examined the dorsal muscle mass of the chicken embryonic forelimb, which undergoes splitting during the same developmental period, and analyzed the spatial relationships among muscles, blood vessels, and nerves (Fig. 4A–D). Before splitting, at E5.0 (HH26), the dorsal muscle mass, similar to the ventral muscle mass, appeared as a single, thin muscle mass (Fig. 4E’, F’). At this stage, neither prominent vascular extension nor nerve projection into the muscle mass was observed (Fig. 4E, E’, F, F’). At E5.5 (HH27), when splitting was in progress, the dorsal muscle mass remained continuous, but a putative splitting region was apparent, as in the ventral muscle mass (white arrowheads in Fig. 4G’, H’). However, unlike the ventral muscle mass, no prominent vascular extension or nerve projection was observed around this region. Instead, vascular extension and nerve projections into the muscle mass were observed at locations distinct from the splitting region (white arrows in Fig. 4G, G’, H, H’). After splitting, at E6.0 (HH28), the dorsal muscle mass had separated into dorso-anterior and dorso-posterior muscle masses (Fig. 4I’, J’). No blood vessels were observed between the two muscle masses, and no prominent vascular extension into either muscle mass was evident (Fig. 4I, I’). Nerve projections were observed within the dorso-anterior muscle mass, whereas no prominent nerve projection was detected within the dorso-posterior muscle mass (Fig. 4J, J’).

**Figure 4.**
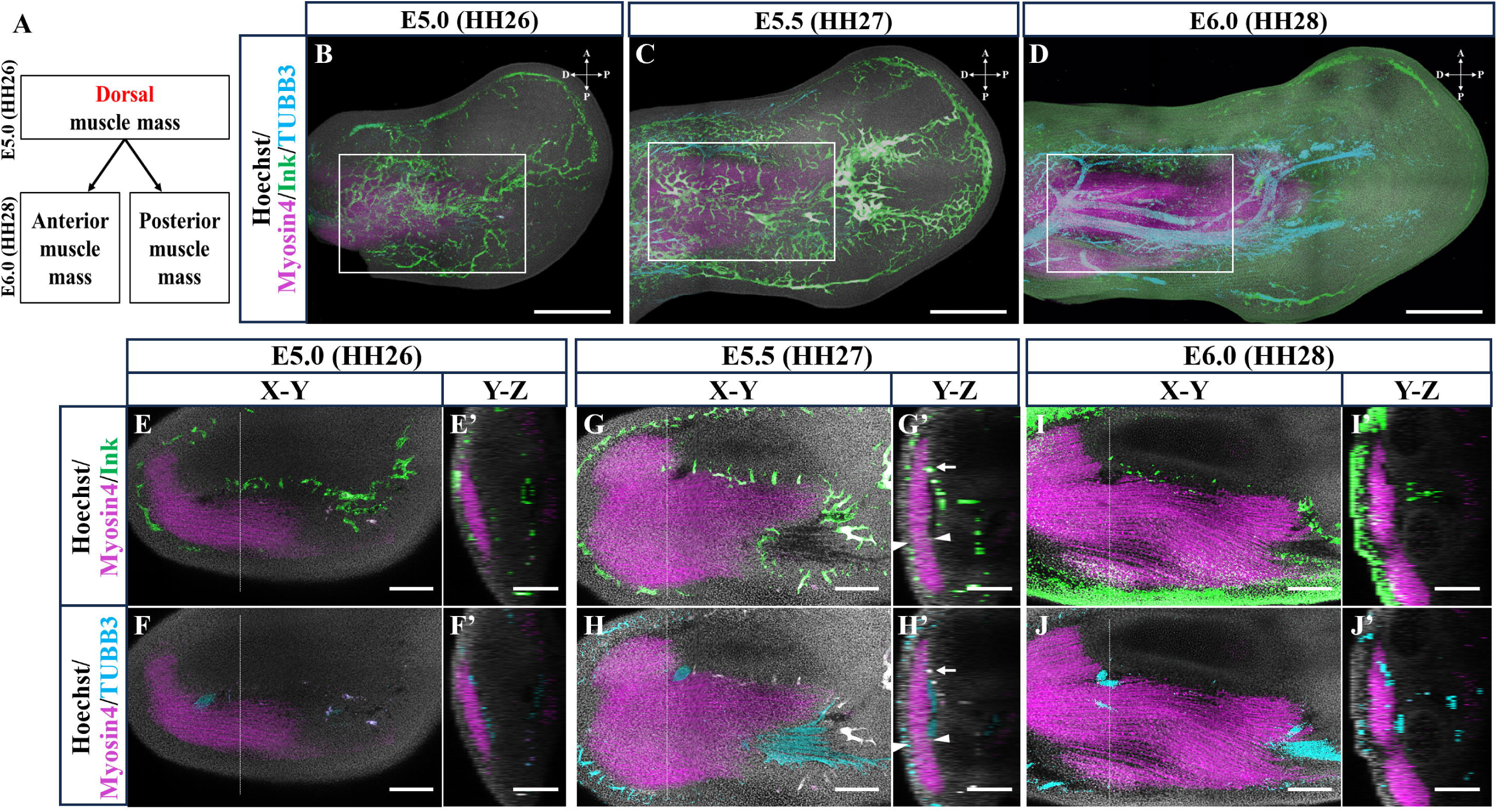
Spatial relationships among muscle, blood vessels, and nerves during splitting of the dorsal muscle mass. (A) Schematic illustration of dorsal muscle mass splitting from E5.0 (HH26) to E6.0 (HH28). (B–D) Muscles (Myosin4), blood vessels (Fluorescent ink; Ink), and nerves (TUBB3) in chicken embryonic forelimbs before splitting (E5.0 [HH26]; B), during splitting (E5.5 [HH27]; C), and after splitting (E6.0 [HH28]; D). White boxes indicate regions shown at higher magnification in E–J’. (E–J’) Higher-magnification X–Y images and corresponding Y–Z views at the positions indicated by dashed lines. At E5.0, no prominent vascular extension or nerve projection into the dorsal muscle mass was observed (E, E’, F, F’). At E5.5, no prominent vascular extension or nerve projection was detected around the putative splitting region (white arrowheads; G, G’, H, H’), whereas vascular extension and nerve projections were observed at locations distinct from the splitting region (white arrows). At E6.0, no blood vessels were observed between the separated dorso-anterior and dorso-posterior muscle masses, and no prominent vascular extension into either mass was evident (I, I’). Nerve projections were present in the dorso-anterior muscle mass but were not prominent in the dorso-posterior muscle mass (J, J’). Nuclei, muscles, blood vessels, and nerves were visualized with Hoechst, anti-Myosin4, fluorescent ink, and anti-TUBB3, respectively. Scale bars: 500 μm in B–D; 200 μm in E–J and E’–J’.

Together, these observations indicate that blood vessels and nerves show distinct spatial patterns during splitting of the dorsal muscle mass and that these patterns differ from those observed in the ventral muscle mass.

## DISCUSSION

In this study, we optimized tissue-clearing conditions compatible with fluorescent vascular labeling to investigate the three-dimensional spatial relationships among muscles, blood vessels, and nerves in the developing chicken embryonic forelimb, with particular emphasis on their relationships during muscle splitting. SeeDB2 preserved fluorescence from all three vascular labels tested—fluorescent ink, LCA-lectin, and the fluorescent lipid tracer PKH26 (Fig. 1). We further established a modified SeeDB2 protocol incorporating blood decolorization, which improved tissue transparency while preserving PKH26 fluorescence (Fig. 2). Using this approach, we analyzed the early splitting process of the dorsal and ventral muscle masses and found that the spatial relationships among muscles, blood vessels, and nerves differed between the two regions. In the ventral muscle mass, vascular extensions were observed around the splitting region at E5.5 (Fig. 3G, G’), whereas no comparable vascular extension was detected around the splitting region of the dorsal muscle mass despite ongoing splitting at the same developmental stage (Fig. 4G, G’). Nerve projections also showed distinct spatial patterns between the dorsal and ventral muscle masses. These findings indicate that local vascular extension toward the splitting site is not a universal morphological feature of muscle splitting and suggest that individual splitting events may proceed within distinct local tissue environments.

### Vascular involvement in muscle splitting and dorsal–ventral differences

The vasculature has long been proposed to contribute to muscle splitting in the developing chick limb. Tozer et al. showed that endothelial cells are positioned at future cleavage sites before clear separation of the ventral muscle mass occurs (Tozer et al., 2007). They further demonstrated that ectopic vascularization induced by VEGFA suppresses muscle formation, whereas local inhibition of vascular development results in muscle fusion, supporting a functional role for blood vessels in muscle splitting (Tozer et al., 2007). Consistent with these findings, we observed multiple vascular extensions around the splitting region of the ventral muscle mass at E5.5 (Fig. 3G, G’), followed by separation into the ventro-anterior and ventro-posterior muscle masses at E6.0. Thus, our three-dimensional observations support the spatial association between vasculature and ventral muscle splitting previously described using two-dimensional histological sections.

In contrast, a different pattern was observed in the dorsal muscle mass. Although splitting was already in progress at E5.5, no prominent vascular extension was detected around the splitting region (Fig. 4G, G’). Moreover, vascular extensions were present at other locations within the same dorsal muscle mass (Fig. 4G, G’), making it difficult to explain this difference simply by a general delay in vascular development on the dorsal side. Although the vascular manipulation experiments of Tozer et al. demonstrated that blood vessels can influence muscle patterning, our morphological observations suggest that local vascular extension toward the splitting site is not a feature shared by all splitting events (Tozer et al., 2007).

These findings further suggest that muscle splitting may not be governed by a single type of tissue interaction, but instead may depend on distinct local patterning environments for individual splitting events. Positional information required for limb muscle patterning is not restricted to muscle progenitor cells themselves but is also provided by the resident limb mesenchyme and muscle connective tissue. In particular, Tcf4-expressing connective tissue cells have been implicated in the pre-patterning and morphogenesis of developing muscles (Huang, 2017; Kardon et al., 2003; Scaal & Marcelle, 2018). Tozer et al. further proposed that endothelial cell-derived PDGFB promotes connective tissue and extracellular matrix formation, thereby excluding myogenic cells from future cleavage sites (Tozer et al., 2007). Comparative developmental studies have likewise suggested that extrinsic cues from surrounding tissues, including connective tissue, tendons, vasculature, and nerves, may contribute to the establishment of limb muscle architecture (Smith-Paredes et al., 2022). Thus, rather than acting as the sole determinant of muscle splitting, blood vessels may function as one component of the local mesenchymal/connective tissue environment in some splitting events, whereas dorsal muscle splitting may be regulated by other local mechanisms that do not involve prominent vascular extension toward the splitting site.

### Neural projections and muscle splitting

Neural projections also showed different spatial patterns between the dorsal and ventral muscle masses. In the ventral muscle mass, some nerve projections into the muscle mass were already evident before splitting at E5.0 (Fig. 3F, F’), whereas no prominent nerve projection was observed directly around the splitting region at E5.5 (Fig. 3H, H’). Similarly, in the dorsal muscle mass, no prominent nerve projection was detected around the splitting region at E5.5 (Fig. 4H, H’). Thus, for the early splitting events examined in this study, our observations do not support a simple model in which nerve fibers directly contribute to the physical separation of the muscle mass.

However, these findings do not exclude a role for nerves in muscle splitting. In a comparative analysis of amniotes, including alligator embryos, Smith-Paredes et al. showed examples in which nerves of the brachial plexus extended between developing muscle divisions during early muscle mass cleavage and proposed that nerves may contribute to muscle separation (Smith-Paredes et al., 2022). They also noted that the correspondence between nerves and muscle divisions was stronger at earlier stages and became less apparent as development progressed, suggesting that different cues may contribute at different stages of muscle splitting (Smith-Paredes et al., 2022). In the avian embryonic forelimb, blood vessels and peripheral nerves show broad spatial correspondence, although the vascular network undergoes extensive remodeling through branching and regression, whereas nerve trajectories are relatively stable (Bates et al., 2002). Therefore, the dorsal–ventral differences in neural projection patterns observed here should be considered not only in terms of whether nerves directly drive muscle splitting, but also in the context of the possibility that muscles, blood vessels, and nerves respond to shared or distinct local patterning cues.

### Future analyses incorporating connective tissue, tendon, and cartilage

The E5-E6 stages examined in this study correspond to a period in which not only muscle formation but also the development of surrounding musculoskeletal tissues, including cartilage primordia and tendons, is actively progressing. Comparative developmental analyses have shown that muscle mass cleavage patterns and the basic morphology of individual muscles are established before stable associations with skeletal elements are formed, suggesting that caution is warranted in regarding the skeleton or cartilage itself as a direct driver of early muscle splitting (Smith-Paredes et al., 2022). In contrast, muscle connective tissue contributes to the pre-patterning of muscle organization, and the integration of muscles, tendons, and skeletal elements becomes increasingly interdependent as development proceeds (Huang, 2017; Scaal & Marcelle, 2018). In our Hoechst-stained specimens, regions with distinct nuclear densities were observed near the muscle masses (Fig. 4I’, J’); however, because cartilage and connective tissue were not identified using specific markers in this study, the histological identity of these regions could not be determined.

Future studies should first directly visualize the distribution of connective tissue around splitting sites using markers associated with muscle connective tissue, such as Tcf4, and subsequently extend the analysis to three-dimensional multiplex imaging of tendons and cartilage primordia positive for SOX9 or COL2A1, where appropriate. In particular, for the dorsal muscle splitting observed here without prominent vascular extension, comparing the spatial distributions of connective tissue, tendons, cartilage, and nerves within the same specimen may provide a more comprehensive understanding of the local tissue environments associated with individual splitting events (Huang, 2017; Smith-Paredes et al., 2022).

### Compatibility between vascular labeling and tissue-clearing methods

A wide variety of tissue-clearing methods based on different principles, including hydrophobic, hydrophilic, and hydrogel-based approaches, have been developed, each with distinct advantages and limitations in clearing efficiency, preservation of tissue morphology, and retention of fluorescent signals and biomolecules (Tian et al., 2021; Ueda et al., 2020). Therefore, selecting a clearing method appropriate for the tissue and labeling strategy is important. In the present study, SeeDB2 relatively well preserved vascular signals from all three labels tested: fluorescent ink, LCA-lectin, and PKH26. In contrast, PKH26 fluorescence was markedly reduced after CUBIC treatment, whereas both fluorescent ink and PKH26 signals were substantially attenuated after Sca*l*eS treatment. Because PKH26 is a fluorescent tracer that incorporates into lipid membranes, effects of the clearing procedures on membrane lipids may have influenced signal retention. Indeed, the compatibility of lipophilic tracers with tissue-clearing methods varies considerably among clearing approaches (Tian et al., 2021).

CUBIC-1A contains 10% Triton X-100, a nonionic detergent commonly used for delipidation, raising the possibility that its strong effects on membrane lipids contributed to the reduction in PKH26 fluorescence (Susaki et al., 2015). Although the concentration of Triton X-100 is lower in Sca*l*eS, the Sca*l*eS reagents used in this study contain methyl-β-cyclodextrin and γ-cyclodextrin. Cyclodextrins can act as delipidation reagents through cholesterol extraction, which may likewise have affected PKH26 signal retention (Hama et al., 2015). In contrast, SeeDB2 is a hydrophilic clearing method characterized by iohexol-based refractive-index matching and saponin treatment (Ke & Imai, 2018). PKH26 fluorescence was also retained after the saponin-based blood decolorization step introduced in our modified SeeDB2 protocol. These observations suggest that, under the conditions examined here, avoiding treatments with strong lipid-removing effects may be advantageous for preserving PKH26-based vascular labeling.

The mechanism underlying the attenuation of fluorescent ink signals after Sca*l*eS treatment remains unclear. Sca*l*eS contains several chemical components, including DMSO, but the present results do not allow us to attribute the signal loss to any specific component. We also observed a marked tendency toward loss of fluorescent ink signals after treatment with 3DISCO, an organic solvent-based clearing method (data not shown). Further systematic evaluation of individual clearing reagents will be required to identify the factors that determine the stability of fluorescent ink signals during tissue clearing.

### Potential applications of the modified SeeDB2 protocol

The modified SeeDB2 protocol developed in this study may be particularly useful when fluorescent labels introduced directly into the vasculature are combined with whole-mount immunostaining in the same specimen. In model organisms for which antibodies against vascular endothelial cells or genetic labeling systems are well established, these approaches provide powerful means of visualizing the vasculature. In non-model organisms, however, suitable antibodies and genetic tools may be limited. Lectin-based vascular labeling is also useful, although lectin-binding properties can vary among vascular types and animal species (Jilani et al., 2003; Laitinen, 1987).

In contrast, introducing fluorescent labeling reagents directly into the vascular lumen does not require species-specific antibodies or genetically encoded labeling systems and may therefore be applicable to a relatively broad range of animal species. Accordingly, tissue-clearing methods that preserve such introduced fluorescent signals while remaining compatible with whole-mount immunostaining may provide a useful approach for examining three-dimensional relationships between blood vessels and surrounding tissues in diverse biological specimens, including those from non-model organisms.

## LIMMITATIONS

This study has several limitations. First, cleared specimens were imaged using confocal laser-scanning microscopy. Under the imaging conditions used in this study, the depth over which multi-channel fluorescence images, including the shorter-wavelength Hoechst signal, could be acquired at sufficient resolution was approximately 200 μm. Consequently, in forelimbs at E5.5 and later stages, it was difficult to image the entire tissue thickness from one side, and the three-dimensional information obtained in this study was therefore limited to portions of the regions analyzed. Confocal microscopy is well suited for high-resolution imaging of localized regions but has limitations in imaging speed because of point scanning and in the acquisition of large tissue volumes. In contrast, light-sheet fluorescence microscopy is better suited for rapid volumetric imaging of thick cleared specimens (Tian et al., 2021). Indeed, three-dimensional analyses of muscle and nerve development in chicken embryonic limbs have been performed using CUBIC clearing combined with light-sheet microscopy (Luxey et al., 2020). Future application of such imaging systems should enable more comprehensive analysis of muscle splitting and the associated vascular and neural patterns throughout the forelimb.

Second, the muscle-splitting process was reconstructed by comparing fixed specimens at different developmental stages, E5.0, E5.5, and E6.0, rather than by directly tracking vascular and neural dynamics within the same embryo. Therefore, we cannot determine whether the vascular extensions observed at E5.5 subsequently regressed or were reorganized into different vascular networks. Because the vascular network in the developing avian forelimb is known to undergo dynamic remodeling through branching and regression (Bates et al., 2002), analyses at more closely spaced developmental stages or live imaging will be required to clarify the temporal changes in blood vessels and nerves associated with muscle splitting.

Third, fluorescence retention after SeeDB2 treatment may depend on the fluorophore used. In our experiments, the eFluor 660 signal used for nerve immunostaining tended to decrease when specimens were maintained in 100% Omnipaque 350 for an extended period. Whether this reduction reflects an intrinsic property of eFluor 660 or depends on antibody or specimen conditions remains unclear. Therefore, when using this fluorophore under the present clearing conditions, imaging as soon as practicable after transfer to 100% Omnipaque 350 may help minimize signal loss.

## SUMMARY

In this study, we established a modified SeeDB2 protocol suitable for three-dimensional analysis combining fluorescent vascular labeling with whole-mount immunostaining and used it to examine the spatial relationships among muscles, blood vessels, and nerves during early muscle splitting in the chicken embryonic forelimb. Vascular extensions were observed around the splitting region of the ventral muscle mass, whereas dorsal muscle splitting progressed without comparable local vascular extension. These findings suggest that vascular extension toward the splitting site is not a universal morphological feature of limb muscle development. Neural projections also showed distinct spatial patterns between the dorsal and ventral muscle masses, further suggesting that muscle splitting may proceed within local patterning environments shaped by interactions with multiple surrounding tissues, including connective tissue, vasculature, and nerves. Future multiplex three-dimensional visualization of these tissues within the same specimen should help clarify the cellular and tissue interactions that contribute to individual muscle-splitting events.

## DATA AVAILABILITY

Serial X–Y and Y–Z image series for the samples analyzed in this study have been deposited in Zenodo and will be made publicly available upon submission of this manuscript.

## SUPPLEMENTAL MATERIAL

Serial X–Y and Y–Z image series are available at Zenodo: https://doi.org/10.5281/zenodo.19650455

## ACKNOWLEDGMENTS

We are grateful to Dr. Tadashi Nomura and Dr. Toshio Takahahsi for valuable comments for this work.

## AUTHOR CONTRIBUTIONS

Y.T. and S.T. designed the study. S.T. performed experiments using chicken embryos, including vascular labeling and sample fixation. Y.T. performed tissue clearing and immunostaining. Y.T. and S.T. performed confocal imaging and data analysis. Both authors wrote the original manuscript and revised it.

## REFERENCES

Bates, D., Taylor, G. I., & Newgreen, D. F. (2002). The pattern of neurovascular development in the forelimb of the quail embryo. Developmental Biology, 249(2), 300–320. 10.1006/dbio.2002.0771

Hama, H., Hioki, H., Namiki, K., Hoshida, T., Kurokawa, H., Ishidate, F., Kaneko, T., Akagi, T., Saito, T., Saido, T., & Miyawaki, A. (2015). Sca*l*eS: An optical clearing palette for biological imaging. Nature Neuroscience, 18(10), 1518–1529. 10.1038/nn.4107

Hamburger, V., & Hamilton, H. L. (1951). A series of normal stages in the development of the chick embryo. Journal of Morphology, 88(1), 49–92. 10.1002/jmor.1050880104

Huang, A. H. (2017). Coordinated development of the limb musculoskeletal system: Tendon and muscle patterning and integration with the skeleton. Developmental Biology, 429(2), 420–428. 10.1016/j.ydbio.2017.03.028

Jilani, S. M., Murphy, T. J., Thai, S. N. M., Eichmann, A., Alva, J. A., & Iruela-Arispe, M. L. (2003). Selective binding of lectins to embryonic chicken vasculature. Journal of Histochemistry & Cytochemistry, 51(5), 597–604. 10.1177/002215540305100505

Kardon, G., Harfe, B. D., & Tabin, C. J. (2003). A Tcf4-positive mesodermal population provides a prepattern for vertebrate limb muscle patterning. Developmental Cell, 5(6), 937–944. 10.1016/S1534-5807(03)00360-5

Ke, M.-T., & Imai, T. (2018). Optical clearing and index matching of tissue samples for high-resolution fluorescence imaging using SeeDB2. Bio-protocol, 8(20), e3046. 10.21769/BioProtoc.3046

Laitinen, L. (1987). Griffonia simplicifolia lectins bind specifically to endothelial cells and some epithelial cells in mouse tissues. The Histochemical Journal, 19(4), 225–234. 10.1007/BF01680633

Luxey, M., Berki, B., Heusermann, W., Fischer, S., & Tschopp, P. (2020). Development of the chick wing and leg neuromuscular systems and their plasticity in response to changes in digit numbers. Developmental Biology, 458(2), 133–140. 10.1016/j.ydbio.2019.10.035

Scaal, M., & Marcelle, C. (2018). Chick muscle development. The International Journal of Developmental Biology, 62(1–3), 127–136. 10.1387/ijdb.170312cm

Smith-Paredes, D., Vergara-Cereghino, M. E., Lord, A., Moses, M. M., Behringer, R. R., & Bhullar, B.-A. S. (2022). Embryonic muscle splitting patterns reveal homologies of amniote forelimb muscles. Nature Ecology & Evolution, 6(5), 604–613. 10.1038/s41559-022-01699-x

Susaki, E. A., Tainaka, K., Perrin, D., Yukinaga, H., Kuno, A., & Ueda, H. R. (2015). Advanced CUBIC protocols for whole-brain and whole-body clearing and imaging. Nature Protocols, 10(11), 1709–1727. 10.1038/nprot.2015.085

Tainaka, K., Kubota, S. I., Suyama, T. Q., Susaki, E. A., Perrin, D., Ukai-Tadenuma, M., Ukai, H., & Ueda, H. R. (2014). Whole-body imaging with single-cell resolution by tissue decolorization. Cell, 159(4), 911–924. 10.1016/j.cell.2014.10.034

Takase, Y., Tadokoro, R., & Takahashi, Y. (2013). Low cost labeling with highlighter ink efficiently visualizes developing blood vessels in avian and mouse embryos. Development, Growth & Differentiation, 55(9), 792–801. 10.1111/dgd.12106

Tian, T., Yang, Z., & Li, X. (2021). Tissue clearing technique: Recent progress and biomedical applications. Journal of Anatomy, 238(2), 489–507. 10.1111/joa.13309

Tozer, S., Bonnin, M.-A., Relaix, F., Di Savino, S., García-Villalba, P., Coumailleau, P., & Duprez, D. (2007). Involvement of vessels and PDGFB in muscle splitting during chick limb development. Development, 134(14), 2579–2591. 10.1242/dev.02867

Ueda, H. R., Ertürk, A., Chung, K., Gradinaru, V., Chédotal, A., Tomancak, P., & Keller, P. J. (2020). Tissue clearing and its applications in neuroscience. Nature Reviews Neuroscience, 21(2), 61–79. 10.1038/s41583-019-0250-1

